# Photochemical NOT Gate for DNA Computing

**DOI:** 10.1101/2020.07.13.201293

**Authors:** Cole Emanuelson, Anirban Bardhan, Alexander Deiters

## Abstract

DNA-based Boolean logic gates (AND, OR and NOT) can be assembled into complex computational circuits that generate an output signal in response to specific patterns of oligonucleotide inputs. However, the fundamental nature of NOT gates, which convert the absence of an input into an output, makes their implementation within DNA-based circuits difficult. Premature execution of a NOT gate before completion of its upstream computation introduces an irreversible error into the circuit. We developed a novel DNA gate design utilizing photocaging groups that prevents gate function until irradiation at a certain time-point. Optical activation provides temporal control over circuit performance by preventing premature computation and is orthogonal to all components of DNA computation devices. Using this approach, we designed NAND and NOR logic gates that respond to synthetic microRNA inputs. We further demonstrate the utility of the NOT gate within multi-layer circuits in response to a specific pattern of four microRNAs.

## Introduction

Owing to its fully programmable interactions, thermodynamic stability, and synthetic accessibility, DNA has become a versatile material for the construction of molecular devices.^1–5^ DNA computation devices seek to model circuits using individual single-stranded DNA molecules as inputs and outputs that are computed through strand displacement cascades within logic gates composed of double-stranded DNA.^6–8^ In the most common implementation, these DNA-based logic gates utilize a series of toehold-mediated strand displacement reactions in which input oligonucleotides hybridize to complementary toehold regions on a DNA logic gate structure facilitating the displacement of an output oligonucleotide.^9,10^ DNA logic gates have been used to construct devices that control protein function,^11^ join split enzymes,^12^ create aptamer-nanoparticle assemblies,^13^ sense metal ions,^14^ and release small molecules^15^ and their assembly into complex molecular circuits can be utilized as neural networks,^16^ DNA nanoprocessors,^17^ surface immobilized DNA walkers,^18^ and cell type recognition.^19^

The use of photochemical methods to control nucleic acid function is a powerful approach due to the ability to regulate light spatially and temporally in a noninvasive manner. A nucleic acid light-control methodology that has been developed by us,^20^ and others,^21^ is the introduction of photocaging groups onto the Watson-Crick face of nucleobases thereby blocking base pairing and nucleic acid hybridization. Using this approach, light-induced cleavage of the caging groups has been used to externally trigger the function of antisense agents,^22^ splice switching oligonucleotides,^23^ triplex-forming oligonucleotides,^24^ antagomirs,^25^ siRNAs,^26^ gRNAs,^27^ hybridization chain reaction initiators,^28^ PCR primers,^29^ and DNA logic gates for *in vitro* and *in cellulo* applications.^30,31^ Photochemical control of DNA hybridization is a precise tool that adds an additional layer of regulation to DNA computation circuits that is insulated from the rest of the circuit mechanisms.

In DNA computation, the digital ON and OFF states of logic gates can be represented by the presence or absence of a DNA molecule. This designation makes the implementation of a NOT gate, which converts the absence of an input into an output, using DNA challenging because premature execution of a NOT gate may result in incorrect output being released (Figure 1A). The DNA strand displacement cascades that represent logic gate operations are a continuous series of reactions that consume each DNA gate as the circuit proceeds. If a gate computes the incorrect output because its input has yet to be produced from an upstream gate, that computation event cannot be undone and the entire circuit becomes unusable. Taking this into account, DNA NOT gates have been implemented using an “inverter” strand to sequester input DNA, which has a limitation as it can only be used in the first layer of a multi-layered circuit.^32^ In a separate DNA logic gate design, which relies on a reversible toehold exchange reaction, Qian *et al*. cleverly developed NOT logic by representing ON and OFF states through separate DNA species, however, this “dual-rail logic” implementation doubles the size and design complexity of any circuit.^33^ Herein, we demonstrate precise temporal control of gate execution to allow the programmable and interchangeable implementation of a NOT gate for DNA computing. By triggering gate execution at a specific timepoint, upstream computation is allowed to complete, thereby ensuring that the correct circuit output is produced. We achieved this by developing a DNA-based NOT gate using photocaging groups that block operation of the gate until light exposure (Figure 1B). Among the existing mechanisms to regulate nucleic acid function, light in many ways represents an ideal external trigger since its application can be readily controlled spatially and temporally.^20,34^ As such, our design utilizes photochemical activation to halt a DNA strand displacement cascade at a specific point, allowing upstream computations to complete first, thereby leading to the correct circuit outcome.

**Figure 1.**
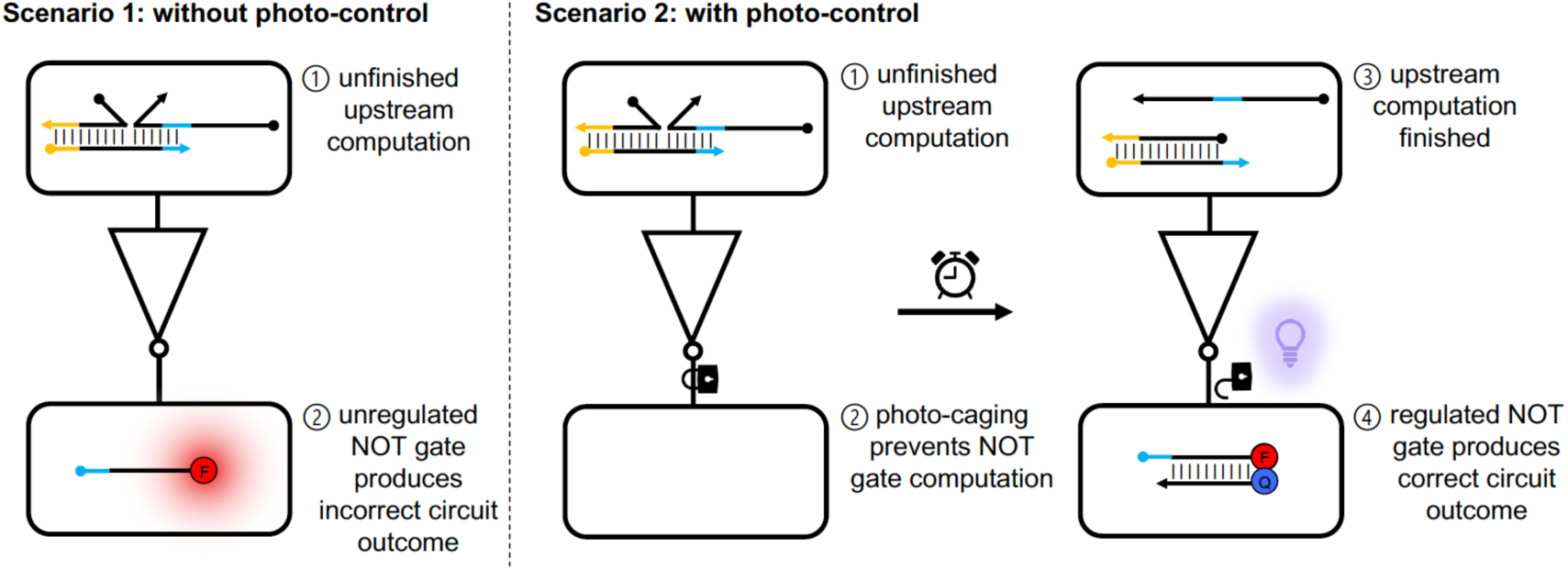
Photo-activated NOT gate principle. In the absence of photo-control, scenario 1, a DNA-based NOT gate will produce an incorrect output because upstream computations have yet to finish and release an input for downstream NOT operation. Photo-control introduces an external trigger, which prevents NOT gate computation in scenario 2 until upstream computations have finished. This additional control allows the correct circuit outcome to be achieved.

## Results and Discussion

To achieve light-triggered NOT gate performance, we designed a double stranded complex that contains two palindromic hairpin-forming domains separated by a loop domain. One of the two NOT gate “arms” is modified with four photocaging groups, while the other is hybridized to a complementary oligo that spans the entire length of the arm. Upon irradiation at 365 nm, the unhybridized arm is decaged, allowing hybridization of the two NOT gate arms yielding a DNA hairpin. This hairpin formation is accompanied by an intramolecular strand displacement of the incumbent NOT gate output, releasing it for downstream computation events (Figure 2A, top). Hairpin formation is inhibited by the hybridization of a complementary NOT gate input to the loop (green) and toehold (blue) domains. Input hybridization prevents hairpin formation after irradiation by sequestering the toehold domain and forming a stable three-stranded complex (Figure 2A, bottom). The four 6-nitropiperonyloxymethylene (NPOM) groups were installed on thymidine nucleotides evenly dispersed in the caged arm of the NOT gate to fully disrupt hybridization and are represented by red squares. While we have chosen to cage thymidine nucleobases here, caging strategies for all four nucleobases have been developed and thus our approach does not introduce any sequence constraints for designing nucleic acid circuits.^20,21^ Four caging groups inhibited hairpin formation, while three or less caging groups did not fully inhibit hairpin formation (data not shown). We have previously shown that the introduction of caging groups can inhibit toehold-mediated strand displacement reactions in a manner proportional to the number of caging groups.^30^ The number of complementary bases adjacent to the incumbent NOT output strand, referred to here as an extended toehold, also influenced hairpin formation. The rate of toehold-mediated strand displacement reaction is dependent on toehold length,^9^ a factor that can be used to limit the rate of DNA circuit processing.^35^ We synthesized NOT gates with zero, one, and three nucleotide-extended toeholds and assessed their purity by gel electrophoresis (Figure S1). With zero extended toehold, hairpin formation and subsequent reporter activation was low. In contrast, a three-nucleotide toehold extension prevented the NOT hairpin formation from being blocked by its input (Figure S2). However, with a one nucleotide toehold, hairpin formation readily occurred in the absence of input upon irradiation and was effectively suppressed by the input. A gel electrophoresis assay confirmed formation of the NOT hairpin and the inhibited NOT gate complex (Figure S3). When combined with a downstream reporter gate, a duplex modified with a fluorophore-quencher pair that is dehybridized upon activation with a matching input DNA, irradiation of the NOT gate in the absence of input DNA triggers the reporter, showing significant (p < 0.001) activation compared to the inhibited gate in the presence of input (Figure 2C). Irradiation times from 1 to 10 min were tested with a minimum irradiation time of 3 min yielding the most activation (Figure S5). Thus, an effective, general, and conditional controlled NOT gate was developed, which can readily be adapted for any input and output DNA sequence.

**Figure 2.**
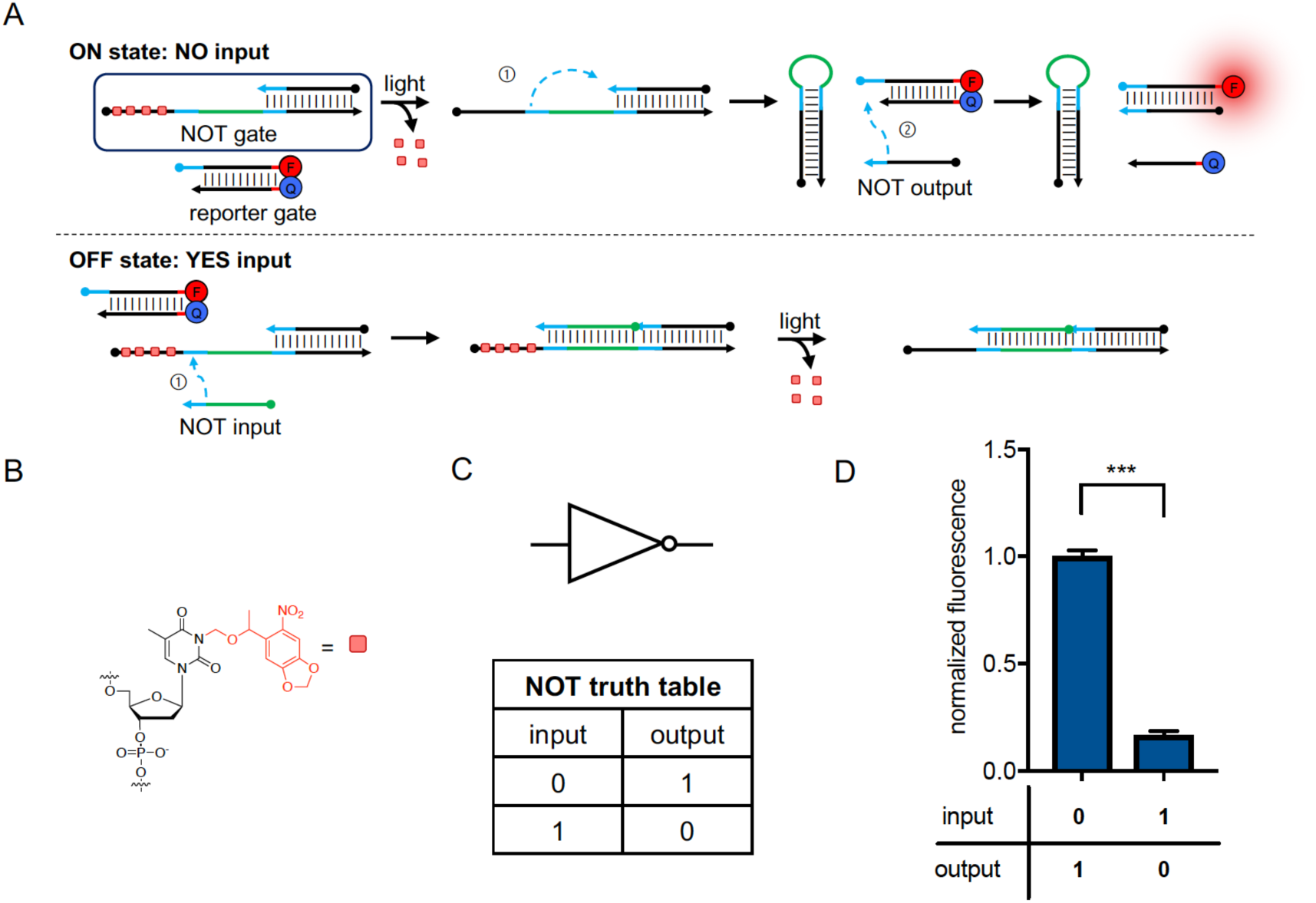
Strand displacement reaction scheme for the photo-activated NOT gate and a corresponding reporter gate. A) In the absence of input (top) light-triggered decaging and gate activation results in intramolecular strand displacement and release of the output strand. The presence of an input strand (bottom) leads to sequestration of the toehold (blue) and loop (green) regions, thereby blocking hairpin formation and preventing intramolecular strand displacement. B) The NPOM (6-nitropiperonyloxymethylene) caging group installed on thymidine nucleotides is represented by a red square. C) NOT gate diagram and corresponding truth table. D) Normalized fluorescence activation of the NOT gate with and without input at 12 h post irradiation. Normalized fluorescence is shown after averaging three independent experiments. Error bars represent standard deviations. ***p<0.001 from unpaired two-tailed Student’s *t*-test. Fluorescence time course data is shown in Supporting Information Figure S4.

Having demonstrated NOT logic, we next tested the universality and modularity of our gate design. As an initial proof-of-principle, we chose to integrate the NOT gate into a circuit containing a simple DNA OR gate. In order to implement OR logic, two translator gates were designed that translate two miRNA input sequences into identical outputs. The output strands released from both translator gates contain a toehold sequence for the loop domain of the NOT gate (Figure 2A). The sequences for the microRNAs miR-486 and miR-126 were adapted to serve as inputs and a specific pattern was chosen where an ON signal represents a non-small cell lung cancer phenotype based on the reported correlation of an absence of specific miRNAs.^36^ The nucleic acid inputs were added to the assembled circuit and toehold mediated strand displacement reactions were allowed to proceed for two hours, followed by optical activation of the NOT gate and completion of the DNA computation cascade. Irradiation at time points earlier than 2 h resulted in reduced inhibition of the NOT gate, demonstrating the importance of time resolved NOT gate activation on circuit performance (Figure S6). Irradiation time points were optimized in a similar manner for subsequent circuit experiments. Overall NOR logic was observed with the detection of either or both miRNA inputs resulting in a significant (p < 0.01) reduction in reporter gate activation compared to the signal in the absence of either input (Figure 2B).

In order to demonstrate that the complete set of Boolean logic functions (AND, OR, and NOT) are compatible with our design, we next combined the NOT gate with an AND translator gate in order to implement NAND logic. As shown in Figure 3B, the combination of both miR-126 and miR-486 inputs is required to generate the NOT gate input strand when the circuit is incubated for 4 h prior to irradiation. Either input alone does not result in the release of the NOT gate input and therefore does not inhibit NOT gate activation and downstream reporter activation (Figure 3B). The level of fluorescence observed for each ON circuit condition, in the absence of input or with either input individually, was not significantly different (p > 0.05), while the fluorescence observed for the OFF circuit condition, in the presence of both inputs, was significantly lower representing the expected NAND gate truth table. Thus, our optically controlled NOT gate can be interfaced with other Boolean logic gates in a simple and modular manner.

**Figure 3.**
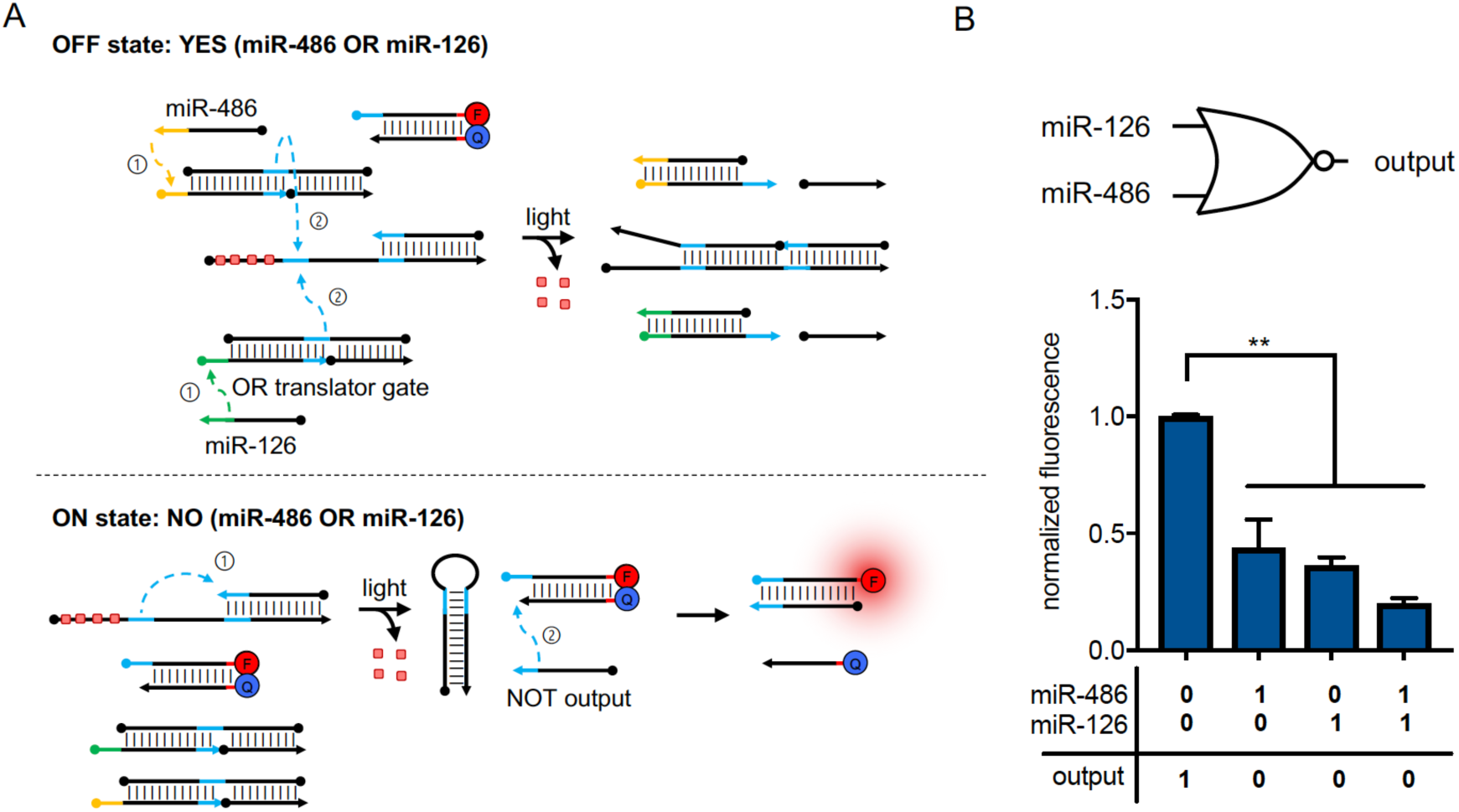
NOR logic gate strand displacement scheme and activation through coupling of an OR gate with the optically triggered NOT gate. A) OR logic is implemented using two translator gates, which are activated by separate miRNAs. Output release by either translator gate (top) suppresses NOT gate activation. In the absence of miRNA input (bottom) NOT gate hairpin formation and downstream reporter activation is triggered by irradiation at 365 nm. B) Quantification of NOR gate activation at 12 h post irradiation. Normalized fluorescence is shown after averaging three independent experiments. Error bars represent standard deviations. **p<0.01 from unpaired two-tailed Student’s *t*-test. Fluorescence time course data is shown in Supporting Information Figure S7.

Having demonstrated the ability to interface the photo-activated NOT gate with individual gates in isolation, we combined the NOR gate components with additional OR and AND gates in order to construct a more complex multi-layered circuit. The combination of four separate logic gates further demonstrates the modularity of the design and its ability to perform within larger logic circuits. The layering of multiple DNA logic gates together has facilitated the construction of more complex systems and has dramatically expanded the functionality of DNA computing devices.^37–39^ The first layer of this multi-layered logic circuit consists of the miR-486 NOR miR-126 gate with a miR-21 OR miR-182 gate, which produce the first and second inputs for the second layer of the circuit consisting of an AND reporter gate. The strand displacement scheme for both ON and OFF cases of the (miR-486 NOR miR-126) AND (miR-21 OR miR-182) circuit is shown in Figure 5A. As shown in Figure 5B, the normalized fluorescence of each of the 3 input conditions corresponding to an ON signal, as indicated within the circuit’s truth table, were not significantly different (p > 0.05). The 13 conditions corresponding to an OFF signal were significantly different from each of the ON conditions (p < 0.01). In conclusion, we successfully demonstrated NOT, OR and AND logic within a multi-layered DNA logic circuit using light to precisely control the cascade of strand displacement reactions and its timing, thereby prevent spurious circuit outcomes.

**Figure 4.**
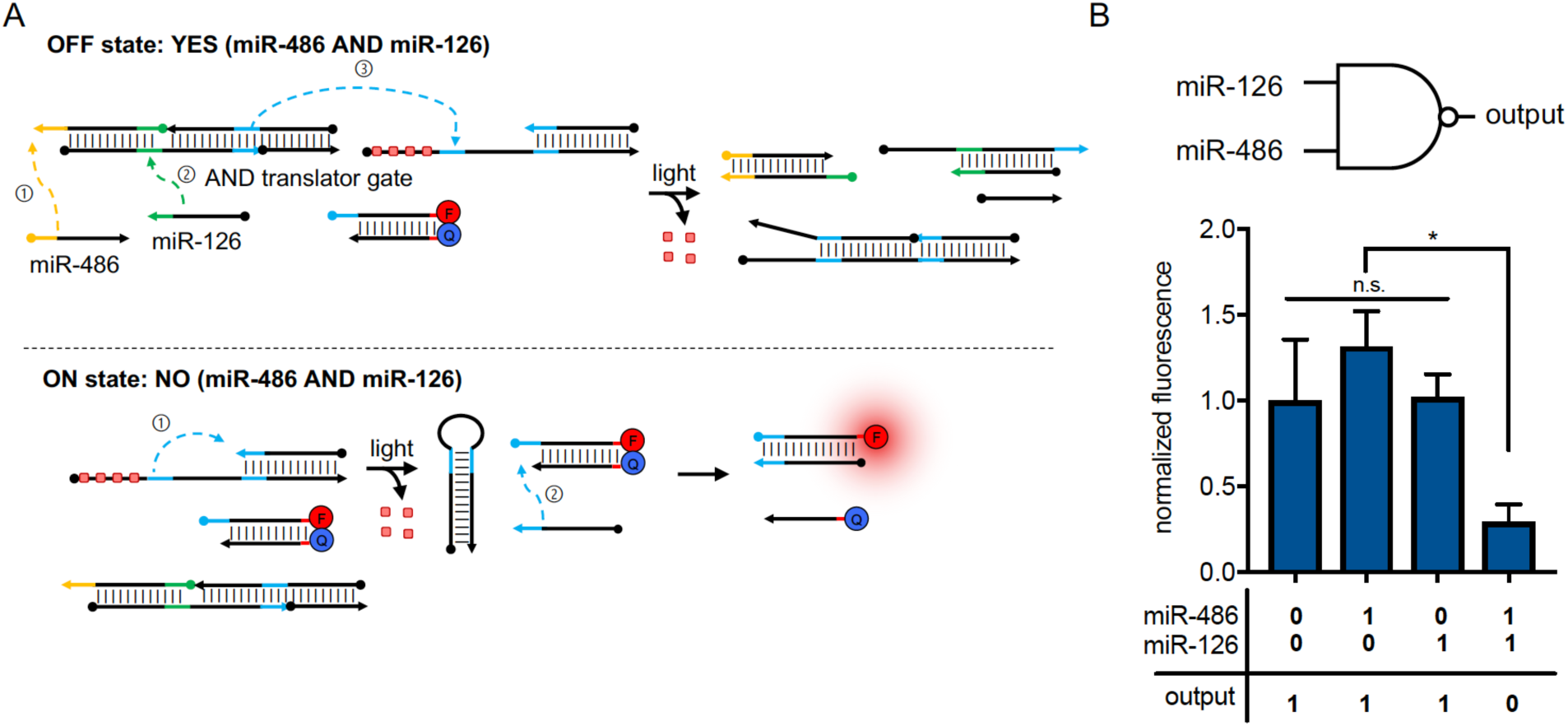
NAND logic gate strand displacement scheme and activation. A) NAND logic is implemented using an AND gate, activated by two miRNA sequences, that is interfaced with the NOT gate and suppresses its function (top). In the absence of the miRNA input (bottom) NOT gate hairpin formation and downstream reporter activation is triggered, leading to a fluorescence output. B) Quantification of NAND gate activation at 12 h post irradiation. Normalized fluorescence is shown after averaging three independent experiments. Error bars represent standard deviations. *p<0.05 from unpaired two-tailed Student’s *t*-test. Fluorescence time course data is shown in Supporting Information Figure S8.

**Figure 5.**
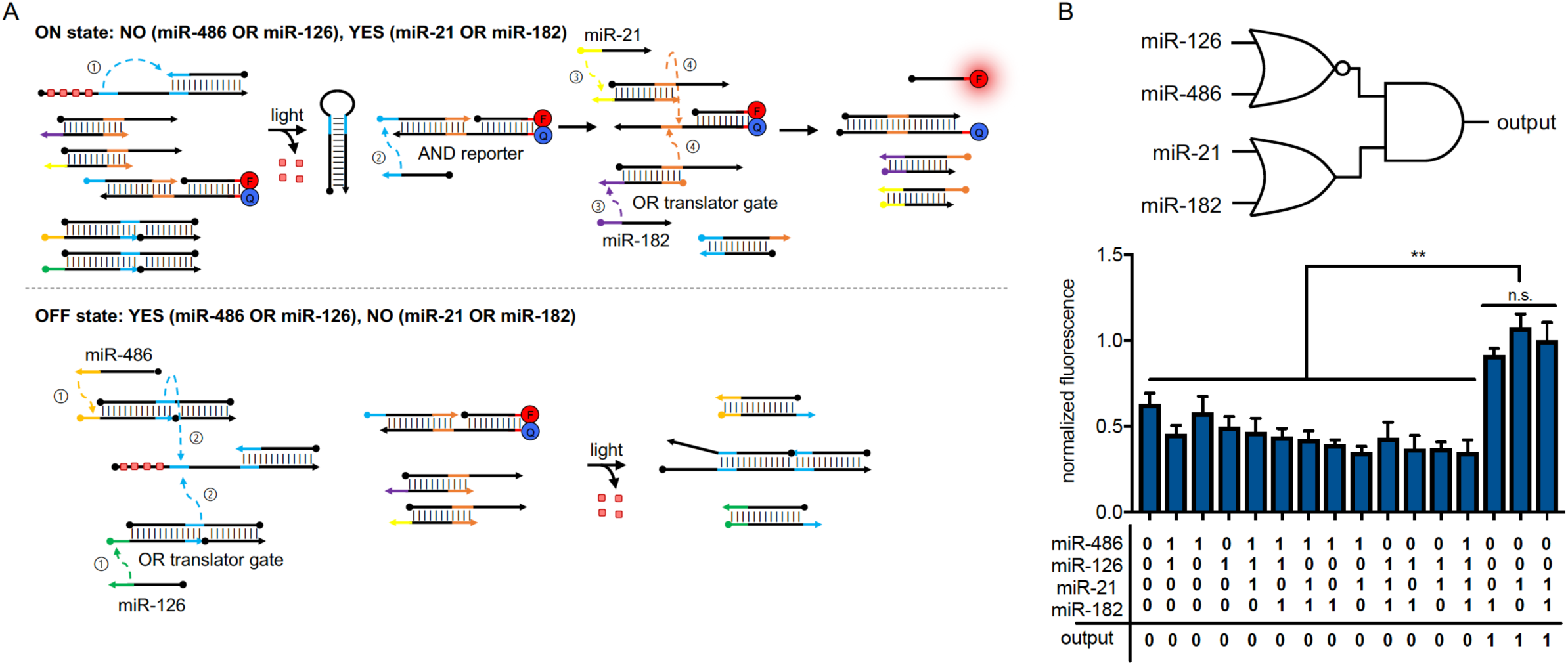
The (miR-486 NOR miR-126) AND (miR-21 OR miR-182) circuit strand displacement scheme and activation. A) In the absence of miR-486 or miR-126 input (top) optical triggering of the NOT gate releases its output, which is the first input to a downstream AND reporter gate. miR-21 OR miR-182 gates are activated by either miR-21 or miR-182 sequences and release the second input for the AND reporter gate. In the presence of miR-126 or miR-486 (bottom) the NOT gate input is generated and prevents hairpin formation and release of the NOT output following irradiation. B) Quantification of (miR-486 NOR miR-126) AND (miR-21 OR miR-182) circuit activation at 20 h post irradiation. Normalized fluorescence is shown after averaging three independent experiments. Error bars represent standard deviations. **p<0.01 from unpaired two-tailed Student’s *t*-test. Fluorescence time course data is shown in Supporting Information Figure S10.

## Conclusions

In summary, a photo-activated NOT gate was designed and synthesized. Precise temporal control over gate activation was demonstrated, following a systematic process of optimizing nucleotide sequence, as well as irradiation and incubation times. Reliable NOT logic was demonstrated in isolation and in conjunction with other DNA gates to implement larger, multi-layer circuits. Previous implementations of DNA NOT gates were limited in applicability^32^ or relied on design rules that increased circuit complexity.^33^ Our photo-activated gate design was successfully integrated within NOT, NOR and NAND circuits, demonstrating its compatibility with a functionally complete set of Boolean logic gates.^40^ Together these gates can be used to implement any Boolean function and will allow for the design of DNA circuits of greater complexity in a modular and straightforward manner. While we have employed a single caging group type, use of orthogonal caging groups which have been used to achieve wavelength-selective oligonucleotide activation^41,42^ along with the development of new caging groups,^43^ will allow for multiplexing of optically activated logic operations within the same circuit. DNA circuitry has been employed for a diverse set of functions including the programming of templated synthesis,^44^ nucleic acid transcription factors,^45^ CRISPR-Cas activity,^46^ microRNA imaging,^47^ ion sensing nanomachines,^48^ and RNA sensing DNA origami.^49^ Moreover, DNA logic gates have been employed in increasingly complex systems including within cellular environments.^31,50–54^ The fast paced evolution of DNA computation and DNA circuitry necessitates the development of new gate designs to complement an increasing number of applications. In addition to general gate function, the sequences of disease-relevant miRNAs were utilized and multi-layered circuits enabled the detection a specific pattern of four separate miRNA sequences, based on a fluorescence readout.

## Experimental Section

### Logic gate preparation and purification

DNA complexes were purified as previously reported.^31^ Briefly, gate duplexes were assembled at 20 μM in 100 μL of 1x TE/Mg^2+^ buffer (Tris-HCl [0.01 M; pH 8.0], EDTA [100 mM], and MgCl_2_ [12.5 mM]) and annealed by cooling the solution from 95 to 12 °C over 30 min in a thermal cycler (Bio-Rad, T100). Detailed descriptions for the assembly of individual gate complexes are described in the supplementary information. Gates were then purified on a 16% native PAGE gel. The full-size duplex bands were identified using a handheld UV lamp (4 W, Analytik Jena, UVL-21) via UV shadowing on a TLC plate, excised, cut into small pieces and eluted overnight in 300 µL TE/Mg^2+^ buffer. For gate complexes with caged oligonucleotides, the purification gel was covered with aluminum foil except for the edge of the lane (approximately 1 cm in width) which was used to locate the band corresponding to the gate complex. Portions of the gel exposed to UV light were not included in the gel extraction. Gate concentrations were determined by UV absorption at 260 nm using Nanodrop ND-1000 (Thermo Fisher Scientific) and calculated with the appropriate duplex extinction coefficient.

### Fluorescence activation measurement

Each reaction was prepared to a 50 µL final volume in 1x TE/Mg^2+^ buffer in a 384-well flat black plate (Greiner). For photo-activation experiments, reactions were incubated in PCR tubes prior to irradiation using a UV-transilluminator (VWR, Dual UV Transilluminator, 80 mW/cm^2^). Following irradiation, reaction mixtures were transferred to an assay plate for fluorescence measurements. TAMRA fluorescence was measured on a TECAN INFINITE M1000 Pro microplate reader (ex/em 545/565 nm) for the indicated time. Fluorescence was normalized to the maximal fluorescence of the ON output condition for each DNA circuit.

### Characterization of strand displacement reactions by native PAGE

The NOT gate was diluted in 1x TE/Mg^2+^ buffer to a final concentration of 50 nM. For conditions with input, input DNA was added to a final concentration of 250 nM. Reaction mixtures were irradiated for 3 min using a UV-transilluminator, where indicated. All reaction mixtures were then separated by gel electrophoresis on a 16% native PAGE gel (140 V, 40 min). For reference, a dsDNA ladder was included (ThermoFisher, cat# SM1233).

## Acknowledgements

This research was supported by the National Science Foundation (CCF-1617041). We thank Nicholas Ankenbruck for help with the design of DNA logic circuits.

## Notes

### Competing Interest Statement

The authors have declared no competing interest.

## References

1 N. Seeman and H. Sleiman, Nat. Rev. Mater., 2017, 3, 1–23.

2 M. Madsen and K. V Gothelf, Chem. Rev., 2019, 119, 6384–6458.

3 F. Hong, F. Zhang, Y. Liu and H. Yan, Chem. Rev., 2017, 117, 12584–12640.

4 J. Bath and A. J. Turberfield, Nat. Nanotechnol., 2007, 2, 275–284.

5 Y. Krishnan and F. C. Simmel, Angew. Chem. - Int. Ed., 2011, 50, 3124–3156.

6 D. Scalise and R. Schulman, Annu. Rev. Biomed. Eng., 2019, 21, 469–493.

7 X. Song and J. Reif, ACS Nano, 2019, 13, 6256–6268.

8 C. Jung and A. D. Ellington, Acc. Chem. Res., 2014, 47, 1825–1835.

9 D. Y. Zhang and E. Winfree, J. Am. Chem. Soc., 2009, 131, 17303–17314.

10 F. C. Simmel, B. Yurke and H. R. Singh, Chem. Rev., 2019, 119, 6326–6369.

11 D. Han, Z. Zhu, C. Wu, L. Peng, L. Zhou, B. Gulbakan, G. Zhu, K. R. Williams and W. Tan, J. Am. Chem. Soc., 2012, 134, 20797–20804.

12 A. Prokup and A. Deiters, Angew. Chem. - Int. Ed., 2014, 53, 13192–13195.

13 M. I. Shukoor, M. O. Altman, D. Han, A. T. Bayrac, I. Ocsoy, Z. Zhu and W. Tan, ACS Appl. Mater. Interfaces, 2012, 4, 3007–3011.

14 S. Bi, Y. Yan, S. Hao and S. Zhang, Angew. Chem. - Int. Ed., 2010, 49, 4438–4442.

15 K. Morihiro, N. Ankenbruck, B. Lukasak and A. Deiters, J. Am. Chem. Soc., 2017, 139, 13909–13915.

16 L. Qian, E. Winfree and J. Bruck, Nature, 2011, 475, 368–372.

17 Y. V. Gerasimova and D. M. Kolpashchikov, Angew. Chem. - Int. Ed., 2016, 55, 10244–10247.

18 C. Jung, P. B. Allen and A. D. Ellington, ACS Nano, 2017, 11, 8047–8054.

19 X. Chang, C. Zhang, C. Lv, Y. Sun, M. Zhang, Y. Zhao, L. Yang, D. Han and W. Tan, J. Am. Chem. Soc., 2019, 141, 12738–12743.

20 Q. Liu and A. Deiters, Acc. Chem. Res., 2014, 47, 45–55.

21 C. Brieke, F. Rohrbach, A. Gottschalk, G. Mayer and A. Heckel, Angew. Chem. - Int. Ed., 2012, 51, 8446–8476.

22 J. M. Govan, R. Uprety, M. Thomas, H. Lusic, M. O. Lively and A. Deiters, ACS Chem. Biol., 2013, 8, 2272–2282.

23 J. Hemphill, Q. Liu, R. Uprety, S. Samanta, M. Tsang, R. L. Juliano and A. Deiters, J. Am. Chem. Soc., 2015, 137, 3656–3662.

24 J. M. Govan, R. Uprety, J. Hemphill, M. O. Lively and A. Deiters, ACS Chem. Biol., 2012, 7, 1247–1256.

25 C. M. Connelly, R. Uprety, J. Hemphill and A. Deiters, Mol. Biosyst., 2012, 8, 2987–2993.

26 V. Mikat and A. Heckel, RNA, 2007, 13, 2341–2347.

27 W. Zhou, W. Brown, A. Bardhan, M. Delaney, A. S. Ilk, R. R. Rauen, S. I. Kahn, M. Tsang and A. Deiters, bioRxiv, 2019, 831974.

28 A. Prokup, J. Hemphill, Q. Liu and A. Deiters, ACS Synth. Biol., 2015, 4, 1064–1069.

29 D. D. Young, W. F. Edwards, H. Lusic, M. O. Lively and A. Deiters, Chem. Commun., 2002, 8, 462–464.

30 A. Prokup, J. Hemphill and A. Deiters, J. Am. Chem. Soc., 2012, 134, 3810–3815.

31 J. Hemphill and A. Deiters, J. Am. Chem. Soc., 2013, 135, 10512–10518.

32 G. Seelig, D. Soloveichik, D. Y. Zhang and E. Winfree, Science (80-.)., 2006, 314, 1585–1588.

33 L. Qian and E. Winfree, Science, 2011, 332, 1193–1196.

34 A. S. Lubbe, W. Szymanski and B. L. Feringa, Chem. Soc. Rev., 2017, 46, 1052–1079.

35 J. Fern, D. Scalise, A. Cangialosi, D. Howie, L. Potters and R. Schulman, ACS Synth. Biol., 2017, 6, 190–193.

36 J. Shen, N. W. Todd, H. Zhang, L. Yu, X. Lingxiao, Y. Mei, M. Guarnera, J. Liao, A. Chou, C. L. Lu, Z. Jiang, H. Fang, R. L. Katz and F. Jiang, Lab. Investig., 2011, 91, 579–587.

37 M. N. Stojanović and D. Stefanović, J. Am. Chem. Soc., 2003, 125, 6673–6676.

38 B. M. Frezza, S. L. Cockroft and M. R. Ghadiri, J. Am. Chem. Soc., 2007, 129, 14875–14879.

39 D. Woods, D. Doty, C. Myhrvold, J. Hui, F. Zhou, P. Yin and E. Winfree, Nature, 2019, 567, 366–372.

40 H. B. Enderton, A Mathematical Introduction to Logic, Harcourt/Academic Press, San Diego, 2nd Ed., 2001.

41 S. Yamazoe, Q. Liu, L. E. McQuade, A. Deiters and J. K. Chen, Angew. Chem. - Int. Ed., 2014, 53, 10114–10118.

42 M. J. Hansen, W. A. Velema, M. M. Lerch, W. Szymanski and B. L. Feringa, Chem. Soc. Rev., 2015, 44, 3358–3377.

43 A. Bardhan and A. Deiters, Curr. Opin. Struct. Biol., 2019, 57, 164–175.

44 W. Meng, R. A. Muscat, M. L. McKee, P. J. Milnes, A. H. El-Sagheer, J. Bath, B. G. Davis, T. Brown, R. K. O’Reilly and A. J. Turberfield, Nat. Chem., 2016, 8, 542–548.

45 L. Y. T. Chou and W. M. Shih, ACS Synth. Biol., 2019, 8, 2558–2565.

46 M. Jin, N. Garreau De Loubresse, Y. Kim, J. Kim and P. Yin, ACS Synth. Biol., 2019, 8, 1583–1589.

47 Z. Cheglakov, T. M. Cronin, C. He and Y. Weizmann, J. Am. Chem. Soc., 2015, 137, 6116–6119.

48 K. H. Leung, K. Chakraborty, A. Saminathan and Y. Krishnan, Nat. Nanotechnol., 2019, 14, 176–183.

49 T. Funck, F. Nicoli, A. Kuzyk and T. Liedl, Angew. Chemie, 2018, 130, 13683–13686.

50 B. Groves, Y.-J. Chen, C. Zurla, S. Pochekailov, J. L. Kirschman, P. J. Santangelo and G. Seelig, Nat. Nanotechnol., 2015, 11, 287–294.

51 G. Chatterjee, Y.-J. Chen and G. Seelig, ACS Synth. Biol., 2018, 7, 2737–2741.

52 Y.-J. Chen, B. Groves, R. A. Muscat and G. Seelig, Nat. Nanotechnol., 2015, 10, 748–760.

53 W. Zhong and J. T. Sczepanski, ACS Sensors, 2019, 4, 566–570.

54 M. Kahan-Hanum, Y. Douek, R. Adar and E. Shapiro, Sci. Rep.,, DOI: 10.1038/srep01535.

