## Supporting Information for "Photochemical NOT Gate for DNA Computing"

### Materials and Methods

#### Oligonucleotide preparation

Unmodified oligonucleotides were purchased from MiliporeSigma (Burlignton, MA), oligonucleotides with fluorophore (3' TAMRA) and quencher (5' Iowa Black Quencher) modifications were purchased from Integrated DNA Technologies (Coralville, IA). Commercial oligonucleotides were diluted to final concentration of 100 µM in nuclease-free water and used directly in logic gate assembly.

The NPOM-caged thymidine phosphoramidite used for synthesizing caged oligonucleotides was prepared following previously established protocols.[1,2] The caged oligonucleotides were synthesized using Expedite 8900 Nucleic Acid Synthesis System, PerSeptive Biosystems, Inc (Framingham, MA) employing standard β-cyanoethyl phosphoramidite chemistry. Syntheses were carried out on a 200 nmol scale using 1000 Å dC derivatized CPG solid phase supports obtained from Glen Research (Sterling, VA). Reagents for automated DNA synthesis were also obtained from Glen Research. Standard synthesis cycles provided by PerSeptive Biosystems were used for all regular nucleobases at 0.067 M concentration while optimized cycles with coupling times of 10 min were applied for the caged phosphoramidite at a 0.05 M concentration. Coupling efficiency in each cycle was monitored by following the release of dimethoxytrityl (DMTr) cations after each deprotection step. No significant reduction in the trityl cation absorbance was noted following the addition of the caged phosphoramidites to the oligonucleotide.

Following solid phase synthesis, cleavage from the support and deprotection of the oligonucleotides was carried out with 1 mL of AMA (ammonium hydroxide/40% aq. methylamine 1:1 v/v) solution at 65 °C for 30 min. After deprotection, the oligonucleotide was concentrated using a Savant OligoPrep OP120 Concentrator, Thermo Fisher Scientific (Waltham, MA) and then analyzed for purity by PAGE (**Figure S1**). The oligonucleotides were directly used in logic gates assembly.

#### Assembly of individual logic gate circuit components

**NOT gate.** The photocaged NOT gate was assembled by adding 20 µL of NOT caged strand (100 µM), 20 µL of NOT output (100 µM), 50 µL of nuclease-free water, and 10 µL of 10x TE/$Mg^{2+}$ buffer to a PCR tube. The reporter gate was assembled by adding 20 µL of fluorophore-labeled strand (100 µM), 20 µL of quencher-labeled strand (100 µM), 50 µL of nuclease-free water, and 10 µL of 10x TE/$Mg^{2+}$ buffer to a PCR tube. Each complex was annealed and purified as described in the Experimental Section of this article. The sequences for each strand used in these gate complexes are shown in **Supporting Table 1**. This procedure was followed for each photocaged NOT gate (NOT-0TH, NOT-1TH, and NOT-3TH:NOT output 22 nt) shown in **Figure S2**, photocaged NOT gate (NOT-1TH:NOT output 22 nt) used in NOT, NOR, and NAND circuits (**Figures 2**, **3**, and **4**) and photocaged NOT gate (NOT-1TH:NOT output 26 nt) used in the (miR-486 NOR miR-126) AND (miR-21 OR miR-182) circuit (**Figure 5**). The NOT gate reporter gate above was used in NOT, NOR, and NAND circuits (**Figures 2**, **3**, and **4**).

**miR-486 NOR miR-126 circuit.** The miR-486 and miR-126 translator gates were assembled separately by adding 20 µL of output strand (100 µM), 20 µL of toehold strand (100 µM), 20 µL of protector strand (100 µM), 30 µL of nuclease-free water, and 10 µL of 10x TE/$Mg^{2+}$ buffer to a PCR tube. The complexes were annealed and purified as described in the Experimental Section of this article. The sequences for each strand used in these gate complexes are shown in **Supporting Table 1**. The circuit diagram is shown in **Figure 3** and **Figure S7**.

**miR-486 NAND miR-126 circuit.** The miR-486 AND miR-126 translator gate was assembled by adding 20 μL of output strand (100 μM), 20 μL of toehold strand (100 μM), 20 μL of backbone strand (100 μM), 20 μL of protector strand (100 μM), 10 μL of nuclease-free water, and 10 μL of 10x TE/$Mg^{2+}$ buffer to a in a PCR tube. The complex was then annealed and purified as described in the Experimental Section of this article. The sequences for each strand used in this gate complex are shown in **Supporting Table 1**. The circuit diagram is shown in **Figure 4** and **Figure S8**.

**(miR-486 NOR miR-126) AND (miR-21 OR miR-182) circuit.** The miR-21 OR miR-182 translator gates were assembled by adding 20 μL of output strand (100 μM), 20 μL of toehold strand (100 μM), 50 μL of nuclease-free water, and 10 μL of 10x TE/$Mg^{2+}$ buffer to a PCR tube. AND reporter gate was assembled by adding 20 μL of toehold strand (100 μM), 20 μL of fluorophore-labeled strand (100 μM), 20 μL of quencher-labeled strand (100 μM), 30 μL of nuclease-free water, and 10 μL of 10x TE/$Mg^{2+}$ buffer to a PCR tube. The complexes were annealed and purified as described in the Experimental Section of this article. The sequences for each strand used in these gate complexes are shown in **Supporting Table 1**. The circuit diagram is shown in **Figure 5** and **Figure S10**.

#### Supporting Figures

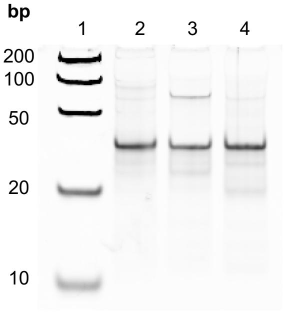

| lane | sample |
|---|---|
| 1 | dsDNA ladder |
| 2 | NOT-caged-strand-0nt-toehold |
| 3 | NOT-caged-strand-1nt-toehold |
| 4 | NOT-caged-strand-3nt-toehold |

**Figure S1.** Gel characterization of the photo activated NOT strands with varying number of hairpin toehold lengths.

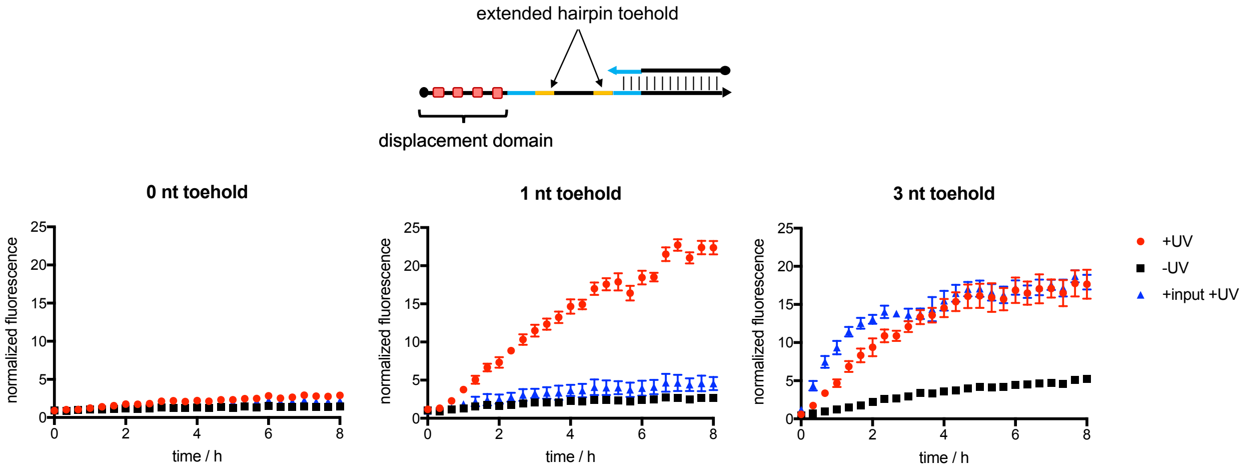

**Figure S2.** Fluorescence activation profile of the photo-activated NOT gate with varying number of hairpin toehold lengths in the presence and absence of input DNA. Data shown are the average fluorescence ± s.d. (n=2) normalized to the average background fluorescence of the reporter gate alone.

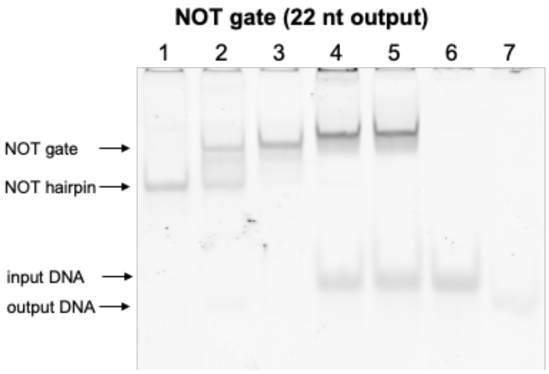

| lane | sample | UV |
|---|---|---|
| 1 | unmodified NOT hairpin | - |
| 2 | NOT gate | + |
| 3 | NOT gate | - |
| 4 | NOT gate + input | + |
| 5 | NOT gate + input | - |
| 6 | input | - |
| 7 | output | - |

**Figure S3.** Gel analysis of the photocaged NOT gate. Irradiation (365 nm, 3 min) results in hairpin formation of the NOT gate (lane 3). Incubation with input DNA prior to irradiation blocks the conversion of the NOT gate to a hairpin (lane 5).

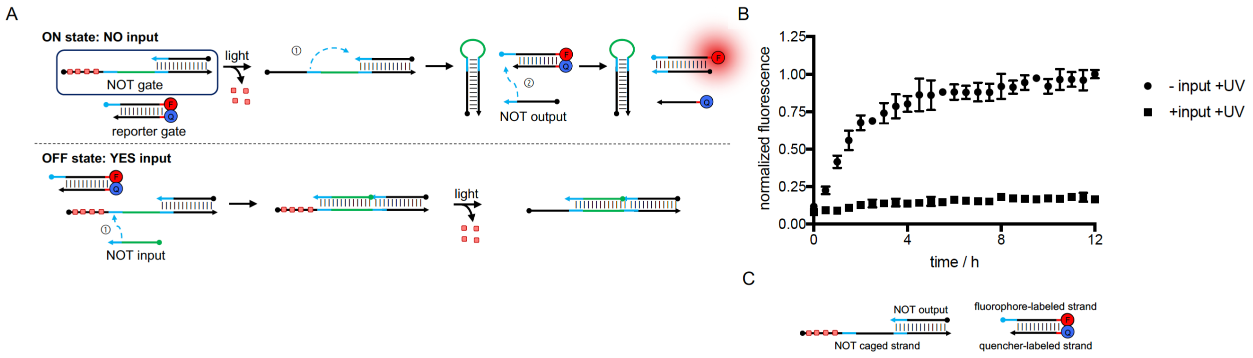

**Figure S4.** Fluorescence time course of the photo-activated NOT gate in the presence and absence of input DNA. A) Strand displacement schematics showing reporter activation and input hybridization to NOT gate. B) Time course activation of reporter gate over 12 h. Bar graphs are shown in **Figure 2**. C) NOT and reporter gate assembly.

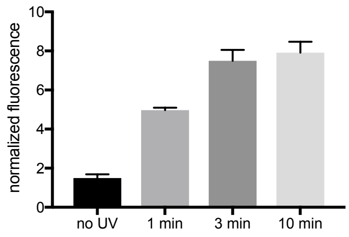

**Figure S5.** NOT gate irradiation time course. NOT gate and reporter gate were irradiated for the indicated time. Average fluorescence is shown at 8 h. Fluorescence was normalized to the average background fluorescence of the reporter gate alone.

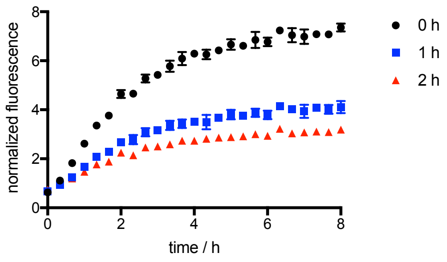

**Figure S6.** miR-126 NOR miR-486 incubation time course. NOT gate, OR gate, and DNA input were incubated for the indicated time prior to irradiation. Fluorescence is normalized to the average background fluorescence of the reporter gate alone.

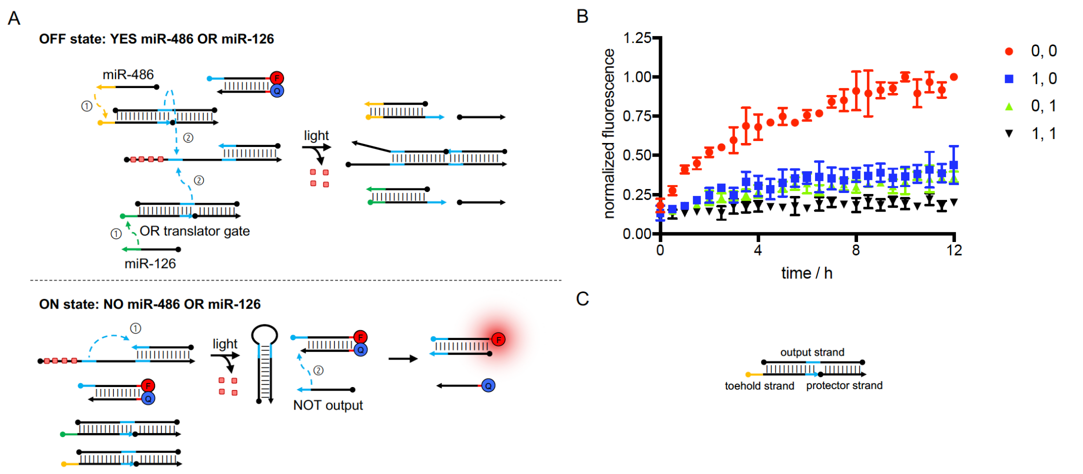

**Figure S7.** Fluorescence activation profile of the photo-activated NOR gate circuit. A) Strand displacement schematic for the miR-486 NOR miR-126 circuit. B) Time course fluorescence activation of reporter gate over 12 h. Average fluorescence of three replicates is shown for input combinations miR-486 only (1, 0), miR-126 only (0, 1) and both miRNAs (1, 1). Bar graphs are shown in **Figure 3**. C) OR translator gate assembly.

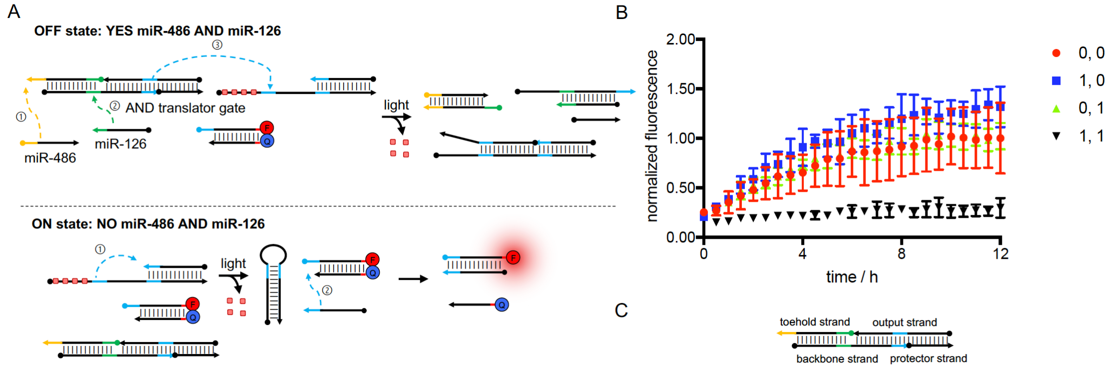

**Figure S8.** Fluorescence activation profile of the photo-activated NAND gate circuit. A) Strand displacement schematic for the miR-486 NAND miR-126 circuit. B) Time course fluorescence activation of reporter gate over 12 h. Average fluorescence of three replicates is shown for input combinations miR-486 only (1, 0), miR-126 only (0, 1) and both miRNAs (1, 1). Bar graphs are shown in **Figure 4**. C) AND translator gate assembly.

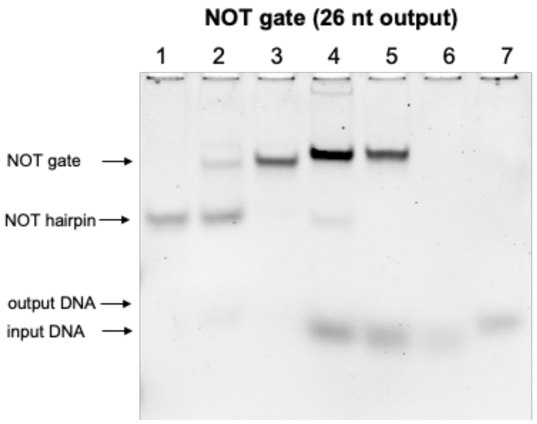

| lane | sample | UV |
|---|---|---|
| 1 | unmodified NOT hairpin | - |
| 2 | NOT gate | + |
| 3 | NOT gate | - |
| 4 | NOT gate + input | + |
| 5 | NOT gate + input | - |
| 6 | input | - |
| 7 | output | - |

**Figure S9.** Gel separation of photocaged NOT gate with 26 nt output DNA used in combination within the (miR-486 NOR miR-126) AND (miR-21 OR miR-182) circuit.

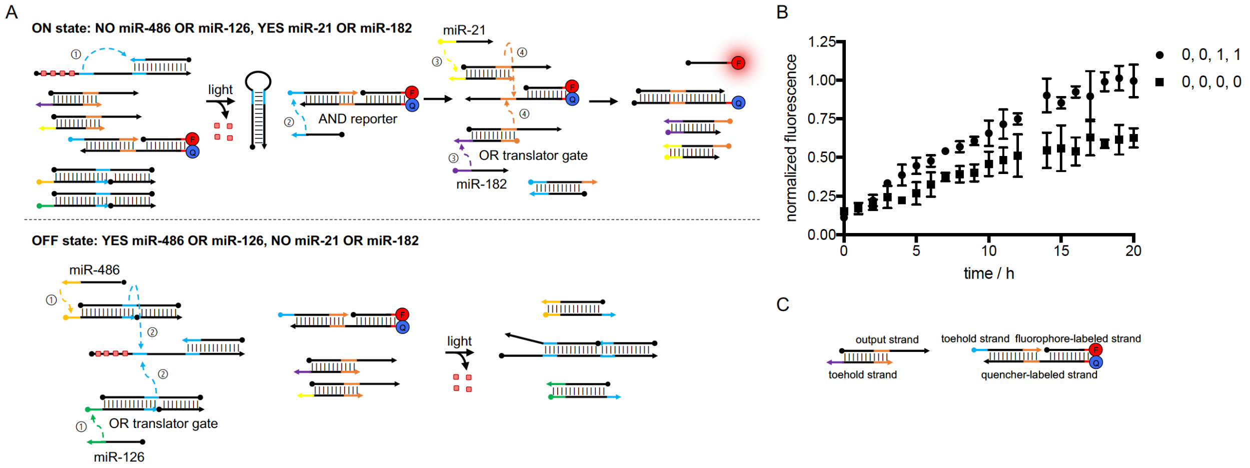

**Figure S10.** Fluorescence activation profile of the photo-activated the (miR-486 NOR miR-126) AND (miR-21 OR miR-182) circuit. A) Strand displacement schematic for the (miR-486 NOR miR-126) AND (miR-21 OR miR-182) circuit. B) Time course fluorescence activation of reporter gate over 20 h. Representative activation profiles for TRUE (0, 0, 1, 1) and FALSE (0, 0, 0, 0) are shown. Bar graphs are shown in **Figure 5**. C) OR translator and AND reporter gate assembly.

#### Supporting Table

**Table S1.** The names and sequences of the oligonucleotides used in this work. Sequences are named based on the complex they are associated with prior to the initiation of a strand displacement cascade. For simplicity, sequences that are used in multiple circuits are shown once. For photocaged NOT gates, the photocaged based are indicated in bold. Fluorophore and quencher modifications are shown based at their point of attachment (5' or 3').

| | | |
|---|---|---|
| | **NOT gate components:** | |
| **NOT gate backbone variants:** | NOT-0TH | G**T**AGC**T**GGT**T**CGAAGCCAG**T**TCTCTGGACTAAC-<br>GAATCGAACTGGCTTCGAACCAGCTAC |
| | NOT-1TH | G**T**AGC**T**GGT**T**CGAA**T**CCAGTTCGACGGACTAACGAAGCGAACTGGAT-<br>TCGAACCAGCTAC |
| | NOT-3TH | G**T**AGC**T**GGT**T**CGAAGCCAG**T**TCGCTGGACTAACGAA-<br>GCGAACTGGCTTCGAACCAGCTAC |
| **NOT gate input and output:** | input strand | GCTTCGTTAGTCCGTCGAACTG |
| | output strand 22 nt | GTAGCTGGTTCGAAGCCAGTTC |
| | output strand 26 nt | CTGCGTAGCTGGTTCGAATACAGTTC |
| | **miR-486 NOR miR-126 circuit components:** | |
| **OR miR-486 translator gate:** | input strand | ACTGTCCTGTACTGAGCTGCCCCGAG |
| | toehold strand | CTCGGGGCAGCTCAGTACAGGACAGTTC |
| | output strand | GCTTCGTTAGTCCGTCGAACTGTCCTGTACTGAGCTGC |
| | protector strand | GACGGACTAACGAAGC |
| **OR miR-126 translator gate:** | input strand | ACTGTCGTACCGTGAGTAATAATGCG |
| | toehold strand | CGCATTATTACTCACGGTACGACAGTTC |
| | output strand | GCTTCGTTAGTCCGTCGAACTGTCGTACCGTGAGTAAT |
| | protector strand | GACGGACTAACGAAGC |

| | | |
|---|---|---|
| **miR-486 NAND miR-126 circuit components:** | | |
| **miR-486 AND miR-126 translator gate:** | miR-486 input strand | TCCTGTACTGAGCTGCCCCGAGCGCA |
| | miR-126 input strand | ACTGTCGTACCGTGAGTAATAATGCG |
| | toehold strand | AATGCGCTCGGGGCAGCTCAGTACAGGA |
| | backbone strand | ACTGAGCTGCCCCGAGCGCATTATTACTCACGGTACGACAGTTC |
| | output strand | GCTTCGTTAGTCCGTCGAACTGTCGTACCGTGAGTAAT |
| | protector strand | GACGGACTAACGAAGC |
| **(miR-486 NOR miR-126) AND (miR-21 OR miR-182) circuit components:** | | |
| **OR miR-21 translator gate:** | input strand | TAGCTTATCAGACTGATGTTGAGCAG |
| | toehold strand | CGCTGCTCAACATCAGTCTGATAAGCTA |
| | output strand | ATCAGACTGATGTTGAGCAGCGAGCGTTCGTCCACCTG |
| **OR miR-126 translator gate:** | input strand | TTTGGCAATGGTAGAACTCACACTGCAG |
| | toehold strand | CGCTGCAGTGTGAGTTCTACCATTGCCAAA |
| | output strand | AATGGTAGAACTCACACTGCAGCGAGCGTTCGTCCACCTG |
| **Reporter gate components:** | | |
| **NOT reporter gate:** | fluorophore-labeled strand | GAACTGGATTCGAACCAGCTACGC/36-TAMSp/ |
| | quencher -labeled strand | /5IAbRQ/GCGTAGCTGGTTCGAATC |
| **NOT AND reporter gate:** | toehold strand | GAACTGGATTCGAACCAGCTACGCAGCG |
| | quencher -labeled strand | /5IABRQ/TGCAGGTGGACGAACGCTCGCTGCGTAGCTGGTTCGAATC |
| | fluorophore-labeled strand | AGCGTTCGTCCACCTGCA/36-TAMSp/ |

#### References

1 H. Lusic and A. Deiters, *Synthesis*, 2006, **2006**, 2147–2150.
2 H. Lusic, D. D. Young, M. O. Lively and A. Deiters, *Org. Lett.*, 2007, **9**, 1903–1906.
3 J. Hemphill and A. Deiters, *J. Am. Chem. Soc.*, 2013, **135**, 10512–10518.
